# Efficient expression of genes in the Drosophila germline using a UAS-promoter free of interference by Hsp70 piRNAs

**DOI:** 10.1101/274589

**Authors:** Steven Z. DeLuca, Allan C. Spradling

## Abstract

Controlling the expression of genes using a binary system involving the yeast GAL4 transcription factor has been a mainstay of *Drosophila melanogaster* developmental genetics for twenty-five years. However, most existing GAL4 expression constructs only function effectively in somatic cells, but not in germ cells during oogenesis, for unknown reasons. A special UAS promoter, UASp was created that does express during oogenesis, but the need to use different constructs for somatic and female germline cells has remained a significant technical limitation. Here we show that the expression problem of UASt and many other Drosophila molecular tools in germline cells is caused by their core *Hsp70* promoter sequences, which are targeted in female germ cells by *Hsp70*-directed piRNAs generated from endogenous *Hsp70* gene sequences. In a genetic background lacking genomic *Hsp70* genes and associated piRNAs, UASt-based constructs function effectively during oogenesis. By reducing *Hsp70* sequences targeted by piRNAs, we created UASz, which functions better than UASp in the germline and like UASt in somatic cells.

## INTRODUCTION

Drosophila is an extremely powerful model organism for studies of animal development and disease because of its low maintenance costs, rapid generation time, and expansive collection of tools to genetically modify its cells. One particularly useful tool is the Gal4/UAS two-component activation system, in which the Gal4 transcriptional activator protein recognizes an upstream activator sequence (UAS) to induce the expression of any gene of interest (Fischer *et al.* 1988; Brand and Perrimon 1993). By controlling the activity of Gal4 with tissue-specific or inducible promoters or the Gal80 inhibitor protein, one can manipulate genes in specific cells or times of development, visualize cell types, probe cell function, or follow cell lineages. One of the most useful applications of these techniques has been to carry out genetic screens by expressing RNAi in targeted tissues or cultured cells (Dietzl *et al.* 2007; Ni *et al.* 2008).

The original pUASt vector from Brand and Perrimon (1993), which contains an *Hsp70*-derived core promoter and SV40 terminator, has undergone several optimizations to improve its expression (Fig 1A). Popular versions, such as the Valium10 or 20 vector used by the Drosophila Transgenic RNAi project (TRiP) (Ni *et al.* 2009; 2011) and the pMF3 vector used by the Vienna Drosophila Research Center (VDRC) GD collection (Dietzl *et al.* 2007) added a ftz intron, and the Janelia Gal4 enhancer project used derivatives of pJFRC81, which added a myosin IV intron (IVS), synthetic 5’UTR sequence (syn21) and viral p10 terminator to boost expression levels across all Drosophila cell types (Figure 1A) (Pfeiffer *et al.* 2012). However, these modifications did not correct UASt‘s major problem-that it drives woefully poor expression in the female germline compared to somatic tissues. Consequently, genetic manipulation in this important tissue has often relied on a special GAL4-activated promoter, UASp, produced by fusing 14 copies of the UAS activator to a germline compatible promoter derived from the P-element, a transposon naturally active in the female germline (Figure 1B) (Rørth 1998). Although UASp expression is qualitatively higher than UASt in the female germline, it is generally known to be lower in somatic tissues.

**Figure 1:**
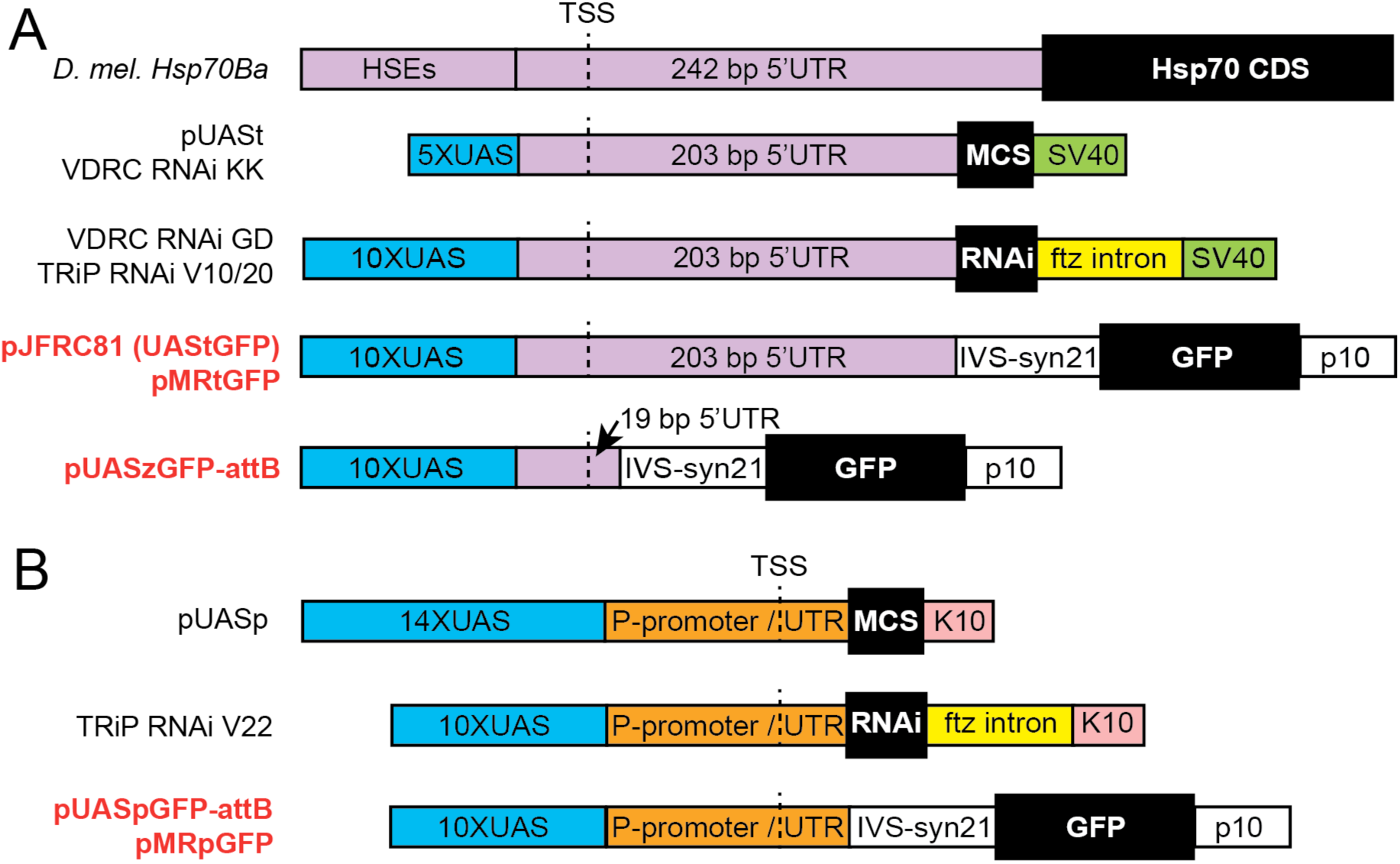
Components of common UAS constructs used by the fly community. (A) Cartoon depicting a Drosophila *Hsp70* gene relative to sequences in UASt-based vectors. In pUASt and VDRC KK lines, multiple copies of optimized Gal4 binding sites (5xUAS) replace heat-inducible enhancers (Heat Shock Elements, HSEs) in a fragment of Hsp70 containing the transcription start site (TSS) and 5’UTR. In derivatives of UASt such as VDRC GD lines and TRiP Valium 10/20 lines, a multiple cloning site (MCS), RNAi constructs, GFP coding sequence, synthetic UTR elements (syn21), and introns (ftz or myosin IV, IVS) replace 39 bp of Hsp70 5’UTR and Hsp70 coding sequence (CDS). Viral-derived SV40 or p10 sequences terminate transcription and contribute to the 3’UTR. For this study, we created a derivative of pJFRC81 (a Janelia-optimized UASt) compatible with MiMIC RMCE (pMRtGFP) as well as pUASz, with a truncated 5’UTR (pUASzGFP-attB). (B) Cartoon depicting the original UASp containing the K10 terminator and P-element promoter, TSS and 5’UTR (in place of the pUASt SV40 terminator and Hsp70 sequences), and the TRiP Valium 22 vector incorporating UASp and a ftz intron (Ni et al. 2011). We created two new UASp vectors, pUASpGFPattB and pMRpGFP, based on pJFRC81and pMRtGFP to directly compare the effect of P-element and Hsp70 sequences on transgene expression. Vector names colored red are used in this study.

The lack of a UAS construct that is widely useful in all Drosophila tissues has remained an obstacle to providing optimum genetic tools to the research community. Transgenic RNAi collections were first constructed using UASt and screening of genes for germline functions has relied on increasing the effectiveness of RNAi by co-expressing Dcr2 or expressing short hairpin RNAi from UASp promoters (Ni *et al.* 2011; Yan *et al.* 2014; Sanchez *et al.* 2016). A significant obstacle to obtaining a widely effective GAL4 vector has been the lack of understanding of the reason UASt functions poorly in germ cells, and the paucity of accurate comparisons between the UASp and UASt promoters in the absence of other significant variables.

## RESULTS AND DISCUSSION

### Difference between UASp and UASt

To study the difference between the UASp and UASt promoters, we first created UASt-GFP and UASp-GFP constructs controlled for other variables between the original UASt and UASp, such as UTR components, introns, terminators, and genomic insertion site. Both constructs were based on pJFRC81 and only varied at the promoter and 5’ UTR of the transcript (Fig 1, red letters). We made these constructs compatible with phiC31-catalyzed recombination-mediated cassette exchange with MiMIC transposons, allowing us to integrate UAS-GFPs into many common sites throughout the genome (Venken *et al.* 2011). Using a previously established protocol (Nagarkar-Jaiswal *et al.* 2015), we recombined both UAS-GFPs into several MiMICS, including MI04106, which resides in a region enriched for ubiquitously expressed genes and active chromatin marks (Filion *et al.* 2010; Kharchenko *et al.* 2011) referred to as “the gooseneck” by Calvin Bridges for its long stretch of low density in salivary gland polytene chromosome preps (Bridges 1935). Consistent with previous reports, UASt drove significantly stronger expression than UASp in all somatic tissues examined while UASp drove significantly stronger expression in the female germline (Fig 2A,B).

**Figure 2:**
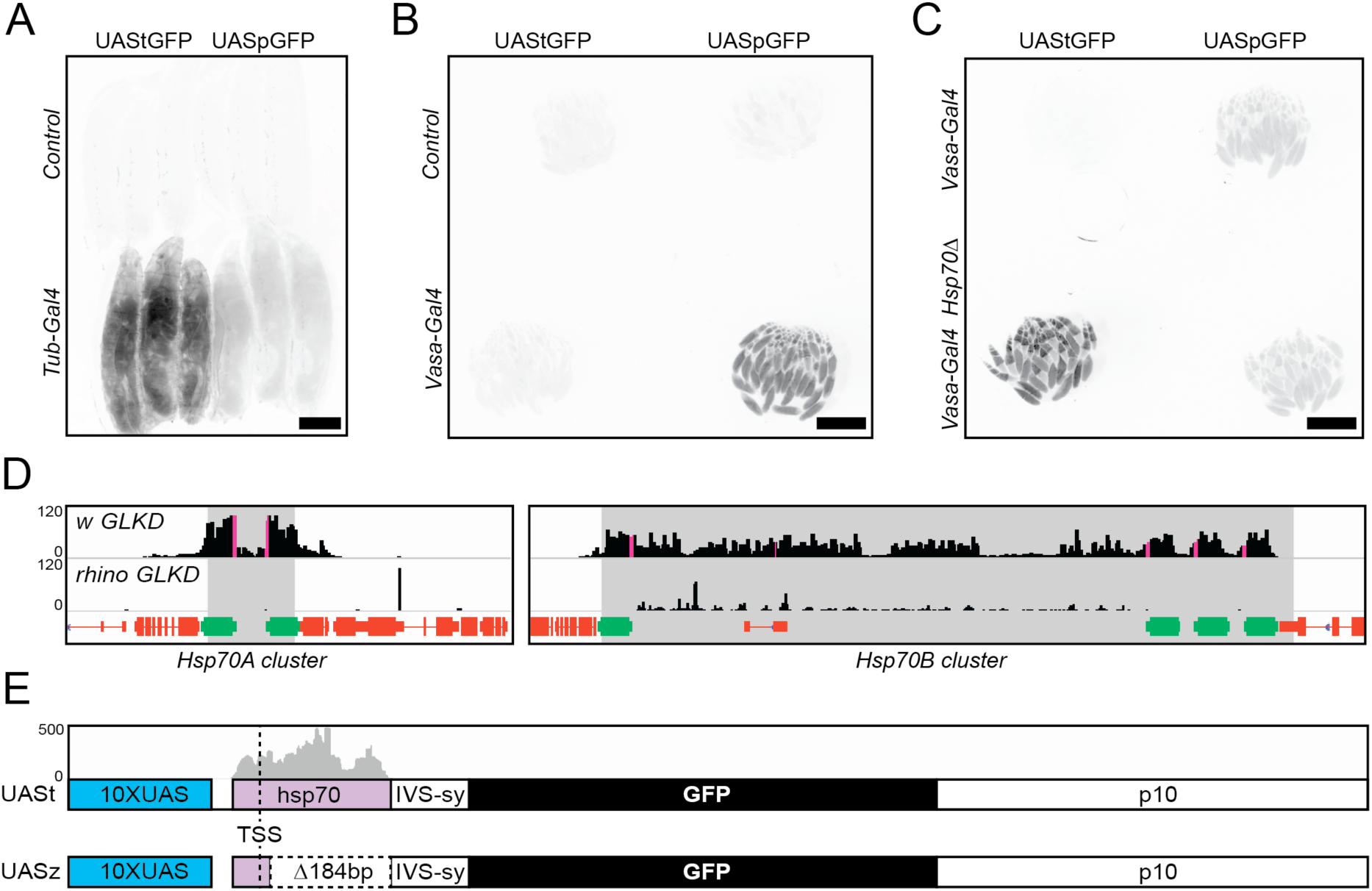
Expression from UASt is greater than UASp in cells lacking *Hsp70* piRNAs. (A-C) pMRtGFP (UAStGFP) and pMRpGFP (UASpGFP) integrated into the same MiMIC site (Mi04106) and crossed to either a control without Gal4 to visualize UAS-GFP leakiness, *Tub-Gal4* for somatic UAS-GFP expression (A), or *Vasa-Gal4* for germline UAS-GFP expression (B,C). Each panel is a single inverted GFP fluorescence image with all 4 genotypes mounted side by side to compare expression levels. Scale bar is 1 mm. (A) Wandering third instar larvae. (B,C) Adult ovaries. (C) Germline UAS-GFP expression in the presence (*Vasa-GAL4*) or absence (*Vasa-GAL4 Hsp70Δ)* of *Hsp70* genes and piRNAs. Image in (B) is longer exposure than (C) to show minimal induction of UAStGFP by vasa-Gal4 in the presence of *Hsp70* piRNAs. (D,E) Genome browser view of whole-ovary-derived piRNAs from (Mohn *et al.* 2014) aligned to *Hsp70A* and *Hsp70B* gene clusters (D), or to pMRtGFP (UAStGFP) (E). (D) piRNA read depth in black. piRNA read depth also mapping to *UASt* in magenta. Control knockdown ovaries (w GLDK) on top. Rhino germline-specific knockdown (rhino GLKD) ovaries on bottom. *Hsp70* genes colored in green. Grey shaded area represents the DNA deleted in the *Hsp70Δ* background. (E) piRNAs mapping to UAStGFP in grey on top. Bottom graphic shows region deleted from UAStGFP to create UASzGFP (Δ184 bp).

### Hsp70 piRNAs repress UASt

We next investigated the reason for the extremely weak UASt expression in the female germline. Several lines of evidence implicated piRNA-directed silencing as a mechanism limiting UASt expression. *Drosophila* piRNAs are ovary and testis-enriched, 23-29 nucleotide (nt) RNAs that complex with Argonaut family proteins and silence transposons through homologous base-pairing-directed mRNA cleavage and heterochromatin formation (Siomi *et al.* 2011). Some of the most successful UASt-based genetic screens in the female germline knocked down piRNA biogenesis genes (Ni et al. 2011; Czech *et al.* 2013; Handler *et al.* 2013). If piRNAs were silencing UASt, then UASt-RNAi against a piRNA biogenesis gene would boost UASt expression leading to maximal knockdown. Where might these UASt-piRNAs originate from? Previously, Mohn et al (2015) characterized an abundance of germline-specific piRNAs mapping to both *Hsp70* gene clusters. Because UASt contains the *Hsp70* promoter and 5’UTR, we hypothesized that germline piRNAs against *Hsp70* may be targeting UASt. When we searched for UASt sequences in the piRNAs identified by Mohn et. al (2015), we identified abundant piRNAs perfectly homologous to UASt (Fig 2D pink bars, and Fig 2E grey bars). Similar to UASt silencing, these UASt piRNAs are restricted to the female germline because germline-specific knockdown of *rhino*, a gene required for *Hsp70* piRNA production eliminates UASt piRNAs from whole ovaries (Fig 2D) (Mohn *et al.* 2014).

To directly test whether *Hsp70* piRNAs silence UASt, we tested UASt expression in *Hsp70Δ* flies (Gong and Golic 2004), which completely lack all genetic loci producing piRNAs homologous to UASt (Fig 2D, grey boxes deleted). Despite missing all copies of the inducible *Hsp70* gene family and related piRNAs, *Hsp70Δ* flies have no significant defects in viability or egg production in the absence of heat stress (Gong and Golic 2006). However, *Hsp70Δ* flies showed greatly enhanced UAStGFP expression. Furthermore, UAStGFP expression was significantly stronger than UASpGFP, which was unaffected by *Hsp70Δ* (Fig 2C). These results argue strongly that UASt is normally silenced by *Hsp70* piRNAs and that UASt is a stronger expression vector than UASp in cells lacking *Hsp70* piRNAs.

### Construction of UASz

We next attempted to create a new version of the UAS expression vector that works well in both the soma and female germline. We hypothesized that eliminating the part of UASt targeted by piRNAs would boost UASt expression by the same amount as eliminating the piRNAS themselves. *Hsp70* piRNAs are homologous to 247 nt of the UASt promoter and 5’UTR. While we could make enough substitutions along this stretch to prevent all possible 23 nt piRNAs from binding, we were afraid this approach might impair important promoter sequences. Instead, we hypothesized that *Hsp70* piRNAs might recognize UASt RNA to initiate piRNA silencing. To prevent *Hsp70* piRNAs from recognizing UASt RNA, we trimmed down the UASt 5’UTR to be shorter than a single piRNA, from 213 nt to 19 nt (Fig 1A, Fig 2E). We named this UTR-shortened UASt variant “UASz,” because we optimistically hoped it would be the last one anyone would make.

### Comparison of UAS vectors

To compare the relative expression levels of our UASz to UASp and UASt, we created all three variants in the same GFP vector backbone (pJFRC81) with a single attB site. We used phiC31 integrase to introduce these UAS-GFP variants into a commonly used genomic site, attP40, and recombined all three inserts with *Hsp70Δ* to determine the influence of *Hsp70* piRNAs on their expression. When combined with *Tub-Gal4*, a somatic Gal4 driver, UASz was expressed at least 4 times higher than UASp in all somatic tissues tested and was equivalent or greater than UASt in some somatic tissues like the larval epidermis and salivary gland (Fig 3A,C,E). However, UASz was expressed at about 40% of UASt in discs, suggesting some elements of the UASt 5’UTR may boost expression in some tissues (Fig 3C,E). To measure germline expression, we crossed the three *UAS-GFPs* to *vasa-Gal4*, which is evenly expressed up to stage 6 of oogenesis. In the germline, UASz was expressed about 4 times higher than UASp at all stages, while UASt was expressed at much lower levels than UASp, except in region 1 of the germarium (Fig 3B,D-F) where piRNA silencing is weaker (Dufourt *et al.* 2014). We conclude that UASz is a superior expression vector to UASp in all tissues, and is equivalent to UASt in many, but not all, somatic tissues.

**Figure 3:**
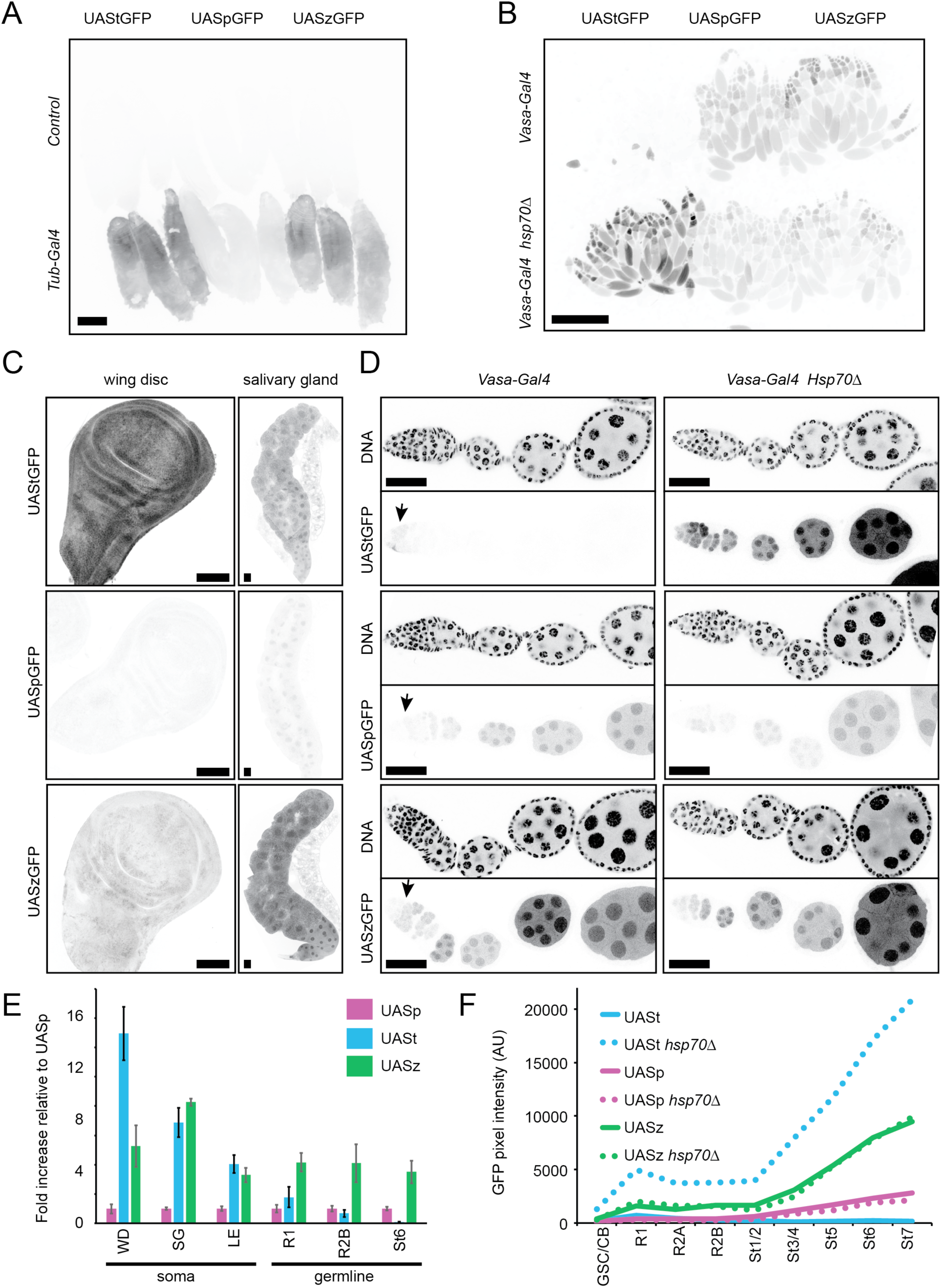
Expression level of *UASz* relative to current *UAS* variants. UAStGFP, UASpGFP, and UASzGFP integrated into a single genomic site, attP40, crossed to control (no *Gal4*) or the indicated *Gal4* driver in either wild type or *Hsp70Δ* background. Inverted GFP fluorescence images of wandering 3^rd^ instar larvae (A), whole adult ovaries (B), or 3^rd^ instar larval wing discs and salivary glands (C). (D) Paired images showing single ovarioles of the indicated genotype imaged in one channel for DAPI (DNA, top) and for GFP fluorescence (bottom). Arrows indicate germline region 1, where piRNA silencing is weakest. Scale bars are 1mm for (A,B) 0.1mm for (C,D). (E) Average GFP fluorescence intensity from UAStGFP and UASzGFP relative to UASpGFP in the indicated tissue expressing *Tub-Gal4* (soma) or *Vasa-gal4* (germline). Error bars indicate standard deviation from the mean from at least 4 samples. (F) Average GFP pixel intensity in germ cells of the indicated stage and genotype. WD = wing disc,SG = salivary gland, LE = larval epidermis, GSC/CB = germline stem cell or cystoblast, R1, R2A, or R2B = germline region 1, 2A, or 2B, St_ = nurse cells of indicated stage number.

Finally, we wanted to test if UASz is still targeted by Hsp70 piRNAs because it contains 63 nt of Hsp70 sequence and about 10% of the putative piRNAs targeting UASt (Fig 2E). We crossed UASzGFP into the *Hsp70Δ* background and compared UASzGFP levels with or without Hsp70 piRNAs. We observed no enhancement of UASzGFP when Hsp70 piRNAs were removed (Fig 3B,D,F). Therefore, Hsp70 piRNAs likely target the UASt but not UASz 5’UTR, consistent with the model that piRNAs must initially recognize RNA but not DNA.

Is UASz the final, fully optimized iteration of a UAS vector? Probably not. UASt without Hsp70 piRNAs induces about twice the expression of UASz in the ovary (Fig 3B,D,F). This twofold advantage of UASt over UASz in the germline or imaginal discs lacking Hsp70 piRNAs is similar to the twofold advantage of UASt over the UAS fused to the Drosophila Synthetic Core Promoter (Pfeiffer *et al.* 2010). Perhaps adding back some sequences within the first 203 nt of the Hsp70 5’UTR while avoiding piRNA recognition may improve UASz.

However, the current iteration of UASz remains an unequivocal upgrade over UASp for all applications and UASz should be preferred over UASt if both germline and soma studies are planned from a single vector. Alternatively, one could boost germline expression of an existing UASt construct by crossing it into the *Hsp70Δ* background.

Current UAS-RNAi collections are heavily biased toward UASt-RNAi-based constructs. To date, the VDRC and DRSC/TRiP RNAi projects used UASt-RNAi to target 12,539 and 8,876 genes, respectively. Germline screens for developmental phenotypes using UASt-RNAi were enriched for phenotypes in germarium region 1 (Yan *et al.* 2014; Sanchez *et al.* 2016), where piRNA silencing is weakest (Dufourt *et al.* 2014) and UASt shows maximum expression (Fig 3D arrow). Perhaps these screens were depleted for developmental defects in later germline stages because of poor UAS-RNAi expression in these stages. Although UASp-RNAi from the Valium22 vector (Fig 1B) increased the efficiency of obtaining phenotypes in a germline screen, only 1,596 genes are currently targeted by this collection (Yan *et al.* 2014). Additionally, when screening somatic cells, Ni et al. (2011) recommend UASt-RNAi because UASp-RNAi gave incomplete knockdowns. Our results revealed that UASp is equally weak in the germline as somatic tissues when compared to UASz (Fig 3E). Therefore, UASp-RNAi may also generate incomplete knockdowns in the germline. To increase germline RNAi expression, we recommend our UASz-RNAi expression vector (Sup Figure 1), which is compatible with previously generated shRNA oligo cloning (Ni *et al.* 2011).

## MATERIALS AND METHODS

### Drosophila strains

*Mef2-Gal4* (BL26882) *w[*]; Kr[If-1]/CyO, P{w+ GAL4-Mef2.R}2, P{w+ UAS-mCD8.mRFP}2 Tub-Gal4* (BL5138) *y[1] w[*]; P{w+ tubP-GAL4}LL7/TM3, Sb[1] Ser[1]*

*FLP/phiC31int* (BL33216) *P{hsFLP}12, y[1] w[*] M{vas-int.B}ZH-2A; S[1]/CyO; Pri[1]/TM6B, Tb[1]*

*Hsp70Δ* (BL8841): *w[1118]; Df(3R)Hsp70A, Df(3R)Hsp70B*

*Vasa-Gal4* was obtained from Zhao Zhang‘s lab: *y[*] w[*];; P{w+ vas-GAL4.2.6}* (Zhao *et al.*2013)

#### New stocks created for this study

Bestgene Inc. introduced pMRtGFP and pMRpGFP into yw flies using a P-transposase helper plasmid and we isolated GFP+ insertions by crossing the F0 to a *Mef2-Gal4* background and scoring for GFP+ muscles. We introduced UAStGFP or UASpGFP into MI04106 and other MiMIC lines using a cross strategy outlined in (Nagarkar-Jaiswal *et al.* 2015). Rainbow transgenics introduced pJFRC81 (UAStGFP-attB), pUASpGFP-attB, and pUASzGFP-attB into attP40 using an X-chromosome encoded phiC31 integrase source and we isolated multiple w+, phiC31 minus insert lines by standard fly genetics.

#### Vectors created for this study

Genescript synthesized pMRtGFP. We created pMRpGFP by replacing the NheI-BglII UASt promoter in pMRtGFP with a SpeI-BglII UASp promoter from Valium22. We created pUASpGFP-attB by replacing the PstI-BglII UASt promoter in pJFRC81 with the PstI-BglII UASp promoter from Valium22. We created UASzGFP-attB by replacing the 259 bp NheI-BglII fragment of pJFRC81 containing the 203 bp *Hsp70* promoter with annealed oligos encoding 63 bp from the 5’ end of the same promoter.

Top oligo: 5’ CTAGCGACGTCGAGCGCCGGAGTATAAATAGAGGCGCTTCGTCTACGGAGCGACAATTCAATTCAAACAAGCAAA 3’

Bottom oligo: 5’ GATCTTTGCTTGTTTGAATTGAATTGTCGCTCCGTAGACGAAGCGCCTCTATTTATACTCCGGCGCTCGACGTCG 3’

We created UASz by replacing the NotI-syn21-GFP-XbaI fragment in UASzGFP with annealed oligos encoding NotI-sny21-BamHI-XhoI-KpnI-SpeI-XbaI

Top oligo: GGCCGCAACTTAAAAAAAAAAATCAAAGGATCCCTCGAGGGTACCACTAGTT

Bottom oligo: CTAGAACTAGTGGTACCCTCGAGGGATCCTTTGATTTTTTTTTTTAAGTTGC

We created UASz1.1 by replacing the KpnI-EcoRI p10 terminator in UASz with a PCR amplified p10 terminator containing Kpn1-XbaI-EcoRI and ApoI tails.

F primer: 5’ CATGGTACCGCCTCTCTAGAGTGTGAATTCTGGCATGAATCGTTTTTAAAATAACAAATCAATTGTTTTATAAT

R primer: 5’ GG<u>AAATTT</u>TCGAATCGCTATCCAAGCCAGCT

We created UASz1.2 by destroying the NheI and EcoRI sites in UASz1.1 by cloning annealed oligos into the NheI-EcoRI backbone.

Top oligo: CTAGGAGCGCCGGAGTATAAATAGAGGCGCTTCGTCTACGGAGCGACAATTCAATTCAAACAAGCAAGATCTGGCCTCGAGT

Bottom oligo: AATTACTCGAGGCCAGATCTTGCTTGTTTGAATTGAATTGTCGCTCCGTAGACGAAGCGCCTCTATTTATACTCCGGCGCTC

To create UASzMiR, we cloned a BglII-XhoI fragment containing the MiR1 cassette and ftz intron from Walium22 into the BglII-XhoI backbone of UASz1.2.

#### Tissue Preparation Imaging and Quantitation

For all experiments, we crossed UAS-GFP or UAS-GFP *Hsp70Δ* males to control (*yw*), *Tub-Gal4/TM3*, homozygous *Vasa-Gal4*, or homozygous *Vasa-Gal4 Hsp70Δ* females. For whole larvae imaging, we picked wandering 3^rd^ instar larvae of various genotypes, aligned them on the same glass slide, and placed them the freezer for 30 minutes prior to imaging. For adult ovary or larval tissue imaging, we fixed dissected tissue with 4% paraformaldehyde for 13 minutes (whole ovary) or 20 minutes (larval tissue) and stained with DAPI in PBS + 0.1% Triton X-100. We imaged GFP fluorescence of semi-frozen whole 3^rd^ instar larvae or whole ovaries mounted in 50% glycerol on a Leica Stereoscope equipped with mercury arc light source, GFP filters, and CCD camera. We imaged GFP fluorescence in larval imaginal discs, salivary glands, and epidermis, and manually separated ovarioles mounted in 50% glycerol using a custom-built spinning disc confocal with 20x 0.8 NA lens. For each genotype and tissue type, we acquired a single plane image from at least 4 individuals using Metamorph software and the same laser power, CCD camera gain, and exposure time between equivalent samples. We measured average pixel intensity in 14 bit images of the GFP channel using Image J. We acquired representative images of single planes through single ovarioles for Figure 2 on a Leica Sp8 scanning confocal with 63x 1.4 NA lens and PMT (for DAPI) and HiD (GFP) detectors using identical settings between samples.

UASt piRNA analysis: We clipped and aligned sequenced small RNA libraries from (Mohn *et al.* 2014) (SRR1187947:control germline knockdown and SRR1187948:rhino germline knockdown) to *D. melanogaster* Genome Release 6 (Hoskins *et al.* 2015) or UAStGFP using the Bowtie2 aligner with no filtering for repetitive mappers (Langmead and Salzberg 2012). We visualized piRNA read depth to UAStGFP or both Hsp70 clusters using the Interactive Genome Browser (Robinson *et al.* 2011).

## ACKNOWLEDGMENTS

We thank members of the Spradling lab for comments. S.Z.D. was a fellow of the Helen Hay Whitney Foundation.

## Supplemental Figure 1

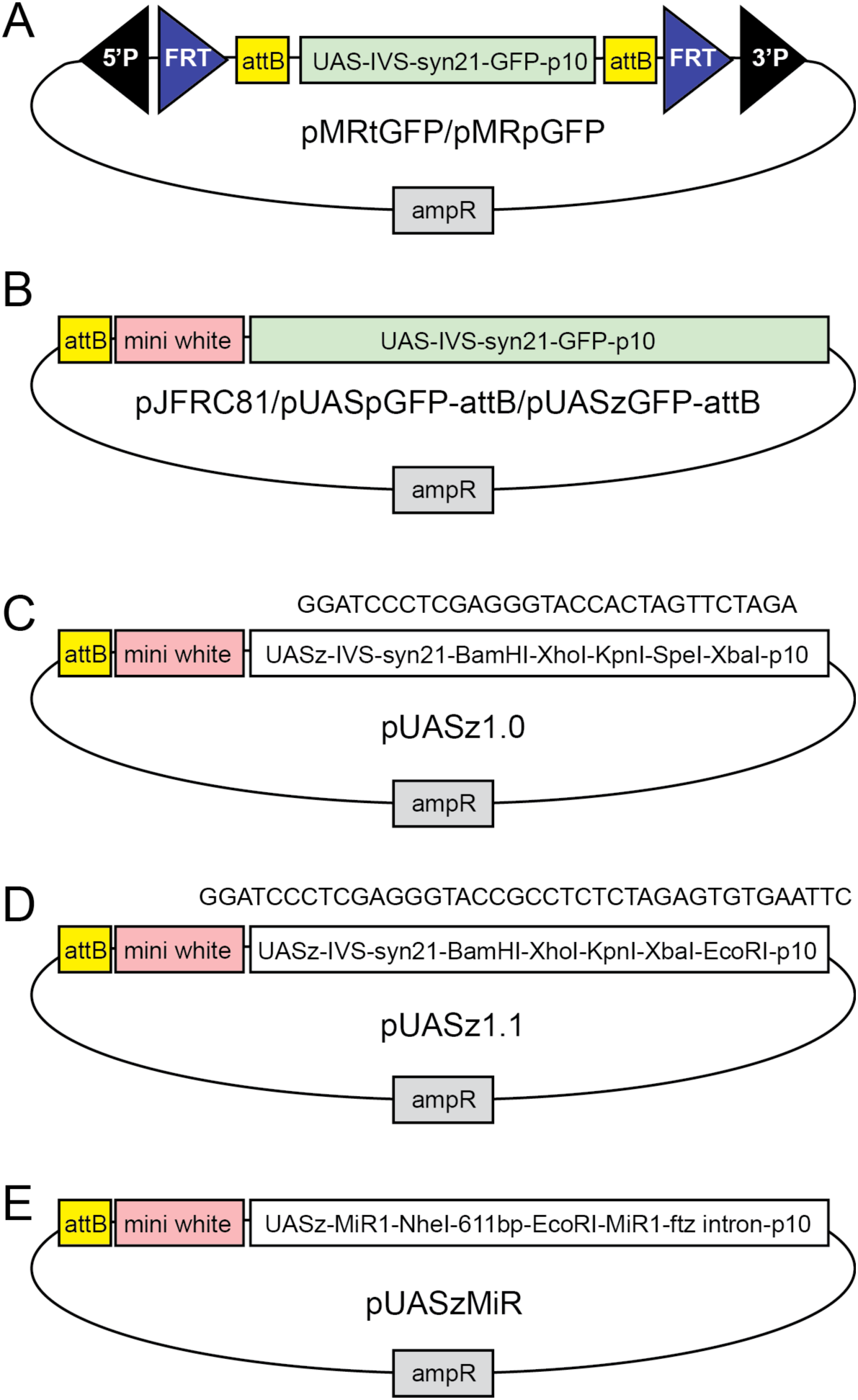
Vectors created for this study (Related to Figure 1). (A) pMRtGFP and pMRpGFP transformation vectors for creating donor flies for in vivo RMCE with MiMICs in the fly genome. The core promoter and 5’UTR of pMRtGFP differs from pMRpGFP as shown in Figure 1. (B) pJFRC81, pUASpGFP-attB, and pUASzGFP-attB mini-white-containing transformation vectors for phi-C31 catalyzed integration into a single attP site in the fly genome. The three plasmids differ at their core promoter and 5’UTR sequences as shown in Figure 1. (C,D) pUASz1.0 and pUASz1.1 are mini-white-containing UASz expression vectors containing slightly different multiple cloning sites shown above each cartoon. (E) UASzMiR is a UASz shRNA expression vector containing the MiR-1 scaffold and ftz intron of Valium22. shRNA encoding oligos can be cloned into the NheI-EcoRI sites as described in Ni et al. (2011).

## REFERENCES

Brand A. H., Perrimon N., 1993 Targeted gene expression as a means of altering cell fates and generating dominant phenotypes. Development 118: 401–415.

Bridges C. B., 1935 SALIVARY CHROMOSOME MAPS With a Key to the Banding of the Chromosomes of Drosophila Melanogaster. J Hered 26: 60–64.

Czech B., Preall J. B., McGinn J., Hannon G. J., 2013 A transcriptome-wide RNAi screen in the Drosophila ovary reveals factors of the germline piRNA pathway. Mol. Cell 50: 749–761.

Dietzl G., Chen D., Schnorrer F., Su K.-C., Barinova Y., Fellner M., Gasser B., Kinsey K., Oppel S., Scheiblauer S., Couto A., Marra V., Keleman K., Dickson B. J., 2007 A genome-wide transgenic RNAi library for conditional gene inactivation in Drosophila. Nature 448: 151–156.

Dufourt J., Dennis C., Boivin A., Gueguen N., Théron E., Goriaux C., Pouchin P., Ronsseray S., Brasset E., Vaury C., 2014 Spatio-temporal requirements for transposable element piRNA-mediated silencing during Drosophila oogenesis. Nucleic Acids Res. 42: 2512–2524.

Filion G. J., van Bemmel J. G., Braunschweig U., Talhout W., Kind J., Ward L. D., Brugman W., de Castro I. J., Kerkhoven R. M., Bussemaker H. J., van Steensel B., 2010 Systematic protein location mapping reveals five principal chromatin types in lDrosophila cells. Cell 143: 212–224.

Fischer J. A., Giniger E., Maniatis T., Ptashne M., 1988 GAL4 activates transcription in Drosophila. Nature 332: 853–856.

Gong W. J., Golic K. G., 2004 Genomic deletions of the Drosophila melanogaster Hsp70 genes. Genetics 168: 1467–1476.

Gong W. J., Golic K. G., 2006 Loss of Hsp70 in Drosophila is pleiotropic, with effects on thermotolerance, recovery from heat shock and neurodegeneration. Genetics 172: 275–286.

Handler D., Meixner K., Pizka M., Lauss K., Schmied C., Gruber F. S., Brennecke J., 2013 The genetic makeup of the Drosophila piRNA pathway. Mol. Cell 50: 762–777.

Hoskins R. A., Carlson J. W., Wan K. H., Park S., Mendez I., Galle S. E., Booth B. W., Pfeiffer B. D., George R. A., Svirskas R., Krzywinski M., Schein J., Accardo M. C., Damia E., Messina G., Méndez-Lago M., de Pablos B., Demakova O. V., Andreyeva E. N., Boldyreva L. V., Marra M., Carvalho A. B., Dimitri P., Villasante A., Zhimulev I. F., Rubin G. M., Karpen G. H., Celniker S. E., 2015 The Release 6 reference sequence of the Drosophila melanogaster genome. Genome Res. 25: 445–458.

Kharchenko P. V., Alekseyenko A. A., Schwartz Y. B., Minoda A., Riddle N. C., Ernst J., Sabo P. J., Larschan E., Gorchakov A. A., Gu T., Linder-Basso D., Plachetka A., Shanower G., Tolstorukov M. Y., Luquette L. J., Xi R., Jung Y. L., Park R. W., Bishop E. P., Canfield T. K., Sandstrom R., Thurman R. E., MacAlpine D. M., Stamatoyannopoulos J. A., Kellis M., Elgin S. C. R., Kuroda M. I., Pirrotta V., Karpen G. H., Park P. J., 2011 Comprehensive analysis of the chromatin landscape in Drosophila melanogaster. Nature 471: 480–485.

Langmead B., Salzberg S. L., 2012 Fast gapped-read alignment with Bowtie 2. Nat. Methods 9: 357–359.

Mohn F., Sienski G., Handler D., Brennecke J., 2014 The rhino-deadlock-cutoff complex licenses noncanonical transcription of dual-strand piRNA clusters in Drosophila. Cell 157: 1364–1379.

Nagarkar-Jaiswal S., DeLuca S. Z., Lee P.-T., Lin W.-W., Pan H., Zuo Z., Lv J., Spradling A. C., Bellen H. J., 2015 A genetic toolkit for tagging intronic MiMIC containing genes. eLife 4: 166.

Ni J.-Q., Liu L.-P., Binari R., Hardy R., Shim H.-S., Cavallaro A., Booker M., Pfeiffer B. D., Markstein M., Wang H., Villalta C., Laverty T. R., Perkins L. A., Perrimon N., 2009 A Drosophila resource of transgenic RNAi lines for neurogenetics. Genetics 182: 1089–1100.

Ni J.-Q., Markstein M., Binari R., Pfeiffer B., Liu L.-P., Villalta C., Booker M., Perkins L., Perrimon N., 2008 Vector and parameters for targeted transgenic RNA interference in Drosophila melanogaster. Nat. Methods 5: 49–51.

Ni J.-Q., Zhou R., Czech B., Liu L.-P., Holderbaum L., Yang-Zhou D., Shim H.-S., Tao R., Handler D., Karpowicz P., Binari R., Booker M., Brennecke J., Perkins L. A., Hannon G. J., Perrimon N., 2011 A genome-scale shRNA resource for transgenic RNAi in Drosophila. Nat. Methods 8: 405–407.

Pfeiffer B. D., Ngo T.-T. B., Hibbard K. L., Murphy C., Jenett A., Truman J. W., Rubin G. M., 2010 Refinement of tools for targeted gene expression in Drosophila. Genetics 186: 735–755.

Pfeiffer B. D., Truman J. W., Rubin G. M., 2012 Using translational enhancers to increase transgene expression in Drosophila. Proc. Natl. Acad. Sci. U.S.A. 109: 6626–6631.

Robinson J. T., Thorvaldsdóttir H., Winckler W., Guttman M., Lander E. S., Getz G., Mesirov J. P., 2011 Integrative genomics viewer. Nat. Biotechnol. 29: 24–26.

Rørth P., 1998 Gal4 in the Drosophila female germline. Mechanisms of Development 78: 113–118.

Sanchez C. G., Teixeira F. K., Czech B., Preall J. B., Zamparini A. L., Seifert J. R. K., Malone C. D., Hannon G. J., Lehmann R., 2016 Regulation of Ribosome Biogenesis and Protein Synthesis Controls Germline Stem Cell Differentiation. Cell Stem Cell 18: 276–290.

Siomi M. C., Sato K., Pezic D., Aravin A. A., 2011 PIWI-interacting small RNAs: the vanguard of genome defence. Nat. Rev. Mol. Cell Biol. 12: 246–258.

Venken K. J. T., Schulze K. L., Haelterman N. A., Pan H., He Y., Evans-Holm M., Carlson J. W., Levis R. W., Spradling A. C., Hoskins R. A., Bellen H. J., 2011 MiMIC: a highly versatile transposon insertion resource for engineering Drosophila melanogaster genes. Nat. Methods 8: 737–743.

Yan D., Neumüller R. A., Buckner M., Ayers K., Li H., Hu Y., Yang-Zhou D., Pan L., Wang X., Kelley C., Vinayagam A., Binari R., Randklev S., Perkins L. A., Xie T., Cooley L., Perrimon N., 2014 A regulatory network of Drosophila germline stem cell self-renewal. Dev. Cell 28: 459–473.

Zhao S., Chen D., Geng Q., Wang Z., 2013 The highly conserved LAMMER/CLK2 protein kinases prevent germ cell overproliferation in Drosophila. Dev. Biol. 376: 163–170.

